# Small molecule activation of the tumor suppressor kinase LKB1

**DOI:** 10.1101/2024.12.17.628051

**Authors:** Lin Song Kretschmer, Dominique C. Mitchell, Jin Liu, LeeAnn Wang, Zheng Wang, Changliang He, Luo Ding, Marc Adler, Rigney E. Turnham, Timothy Kellett, Aidan Keith, Yeonjoo C. Hwang, Gary K. L. Chan, Weicheng Li, Roopa Ramamoorthi, Sarah Lively, Robert A. Drakas, Vijay Ramani, Tingting Qing, Kliment A. Verba, Jonathan M. L. Ostrem, Richard T. Beresis, John D. Gordan

## Abstract

The ability to identify and target oncogenic signals has transformed clinical oncology. Drug development for targeted therapies has historically focused on the inhibition of oncogenic kinases and GTPases. However, many cancer patients do not benefit from targeted approaches because their tumors lack targetable mutations. Therapeutic augmentation of tumor suppressive signaling could be a viable alternative but poses challenges. Specifically, designing compounds capable of stimulating kinase activity is more structurally challenging than inhibitor design, and most kinases lack targetable allosteric pockets. Inactivation of the liver kinase B1 (LKB1) tumor suppressor kinase is associated with poor prognosis and therapeutic resistance. Thus, augmented LKB1 function could be beneficial for cancer patients whose tumors retain intact copies of the gene. LKB1 signals as part of an obligate trimer including the scaffolding protein Mouse protein-25 (MO25) and the pseudokinase (PSK) STE20-related kinase adapter protein (STRAD). As STRAD binds but does not metabolize ATP, it provides the opportunity for a novel activation strategy. We have developed STRAD-binding compounds capable of activating LKB1 and demonstrate the therapeutic benefits of LKB1 activation in a target-dependent manner within cancer cell lines.

## Introduction

Uncontrolled growth in cancer results from dysregulation of oncogenic and/or tumor suppressive signaling. Tumor suppressor kinases preserve homeostasis in normal cells by maintaining polarity, controlling metabolism and controlling responses to inflammation and DNA damage^1–3^. Loss of function of a tumor suppressor kinase is seen in 11% of specimens in The Cancer Genome Atlas (TCGA), with the most commonly inactivated kinases being LKB1 (also known as serine/threonine kinase 11, STK11), MAP2K4 and ATM^4^. Disruption of these pathways often results in synthetic lethal sensitivity to agents targeting parallel pathways^1–3^. Although the altered cell states resulting from inactivation of tumor suppressive kinases have been well studied, their regulation and impact in transformed cells is less well understood. *LKB1* inactivation is observed in 16% of lung adenocarcinomas, 6% of cervical squamous carcinomas, and 1-3% of common malignancies such as breast, ovarian, pancreatic, gastro/esophageal and melanoma in TCGA, with LKB1 inactivation also associated with worse prognosis for both recurrence and overall survival^5,6^. Mouse models and clinical trials have demonstrated the impact of LKB1 on tumor behavior: *Lkb1* loss accelerates tumor initiation in lung and other cancers and is associated with higher rates of metastasis^7,8^. *Lkb1*-deficient mouse cancer models show resistance to chemotherapy and targeted therapy^9^, and clinical studies have connected *LKB1* loss with resistance to KRAS inhibition, EGFR inhibition, chemotherapy, radiotherapy and the PD1 inhibitor nivolumab^10–19^.

Importantly, recent data demonstrate that restoration of LKB1 following its mutational inactivation in *Kras*-driven murine lung cancer models results in dramatically decreasing lung tumor burden^20^. Thus, *LKB1* loss is likely important for both tumor initiation and maintenance, and augmented LKB1 signaling may have tumor suppressive effects in advanced cancers.

LKB1 signals upstream of AMP-activated kinase (AMPK)^21–23^ and 13 other AMPK-related kinases (ARKs)^24^ to regulate protein translation^25^, autophagy^26^, chromatin, cellular differentiation^27–29^, mitochondrial quality control^30,31^, and the cell-intrinsic immune response^32^. Genetic screens have tied these effectors to context-specific cellular outcomes, including a role in pancreatic cancer for the MAP / microtubule affinity-regulating kinases (MARK1-4) regulating the Hippo kinase pathway, which inhibits the YAP/TAZ co-activators to reduce cellular proliferation^33^. Recent studies in cancer cell lines and mouse models revealed that the Salt-inducible kinases (SIK1-3) are the essential effectors of LKB1-mediated tumor suppression *KRAS*-mutant lung cancer, with effects *in vivo* on inflammation and metabolism^7,8^.

While the upstream regulation of LKB1 is still being explored^34^, the biochemistry of the active LKB1 complex allows for development of small molecules targeting its allosteric regulation. LKB1 signals in an obligate trimer that includes the pseudokinase (PSK) STRAD and the adaptor protein MO25. STRAD contacts are necessary for LKB1 activity, as shown by disease-associated LKB1 mutations that disrupt the LKB1/STRAD interface^35,36^. Like most PSKs, STRAD binds and is stabilized by ATP but it does not show phosphotransfer activity^37,38^. PSKs can regulate signaling allosterically, either *in trans* as seen for STRAD and HER3 or *in cis* as in the EIF2AK4 and JAK1/2 kinases which contain tandem PSK and kinase domains^39–41^.

We hypothesized that small molecules targeting the STRAD ATP-binding pocket could activate LKB1 and thus inhibit tumor cell proliferation. Here, we describe compounds termed **S**mall **M**olecule **A**ctivators of **L**KB1 via **S**TRAD (SMALS). We demonstrate the impact of these molecules *in vitro* and in cell models to activate LKB1 and its downstream effectors. These activating tool compounds illuminate the mechanism of action and show the potential of LKB1 activators as a new strategy in cancer treatment.

## Results

### Small molecule binding to STRAD can activate LKB1

First, we asked whether small molecules that bind the pseudokinase STRAD’s ATP-binding pocket could enhance LKB1’s tumor suppressor signaling activity (Fig. 1A). To identify such molecules, we screened a library of 429 advanced compounds for displacement of the fluorescent ATP analog 2’,3’-O-(2,4,7-trinitrocyclohexadienylidene) adenosine 5’triphosphate (TNP-ATP) from the STRAD-MO25 dimer (Fig. 1B). We identified 5 structurally diverse compounds able to reduce TNP-ATP occupancy by greater than 50% at 50 µM. Notably, these included 3 insulin-like growth factor 1 receptor (IGF-1R) inhibitors (NVP-AEW541, AZD3463, and LDK378), as well as the Src family kinase inhibitor saracatinib and the ERK5 inhibitor ERK5-IN-1 (Table. S1). NVP-AEW541 showed the greatest effect on TNP-ATP binding and was confirmed to displace ATP from STRAD with an IC50 of 2.4 μM (Fig. 1C). These 5 compounds were then assessed with a Caliper-based kinase assay using the recombinant LKB1/STRAD/MO25 trimer^42^, with only the pyrrolopyrimidine (PYP) NVP-AEW541 able to directly stimulate LKB1 kinase activity *in vitro* (Fig. 1D).

**Figure 1.**
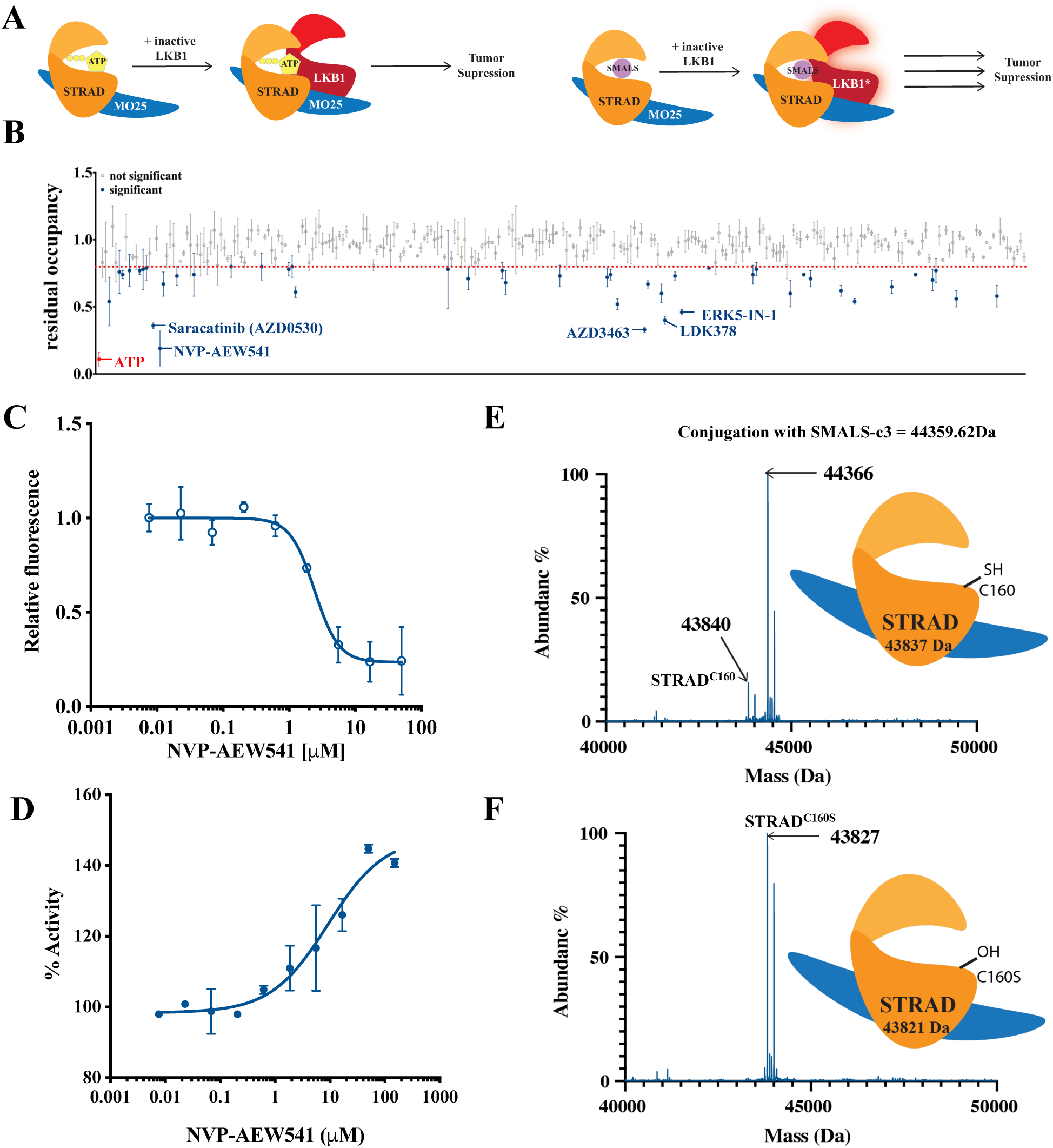
TNP-ATP displacement screen identifies compounds that bind to STRAD and activate LKB1. A. Model of hypothesized enhancement of LKB1 activation through the use of SMALS to stabilize the STRAD pseudokinase “active” conformation, thereby enhancing trimeric complex formation and LKB1 kinase activity. B. TNP-ATP screen of advanced kinase inhibitor library, measuring ability of small molecules to bind STRAD’s ATP-binding pocket. Screen was performed on recombinant STRAD/MO25. Significant hits are colored blue, the top five hits are labeled. ATP is shown and labeled in red. C. TNP-ATP assay validating NVP-AEW541 displacement of ATP from recombinant STRAD/MO25. D. NVP-AEW541 activates LKB1 in an *in vitro* kinase assay using the LKB1/STRAD/MO25 trimer. Kinase activity was read out with a Caliper device. E. Schematic of the calculated size of wild-type of STRAD and its conjugation with the SMALS-c3, anticipated to occur at STRAD Cysteine 160 (C160); LC-MS analysis shows that the majority of wild type of STRAD is conjugated with SMALS-c3 after 6 hours’ incubation. F._C160S_ LC-MS of STRAD incubated with SMALS-c3 demonstrating loss of covalent binding.

To confirm specific binding and develop a tool compound for future studies, we modified NVP-AEW541 by adding a covalent warhead targeting the cysteine residue STRAD^C160^. Liquid chromatography-mass spectrometry (LC-MS) analysis confirmed the conjugation of SMALS-c3 to STRAD-MO25 (Fig.1 E). This binding was lost in a STRAD^C160S^ mutant (Fig.1F), consistent with SMALS binding the ATP pocket on STRAD and supporting our model that SMALS engagement of STRAD ATP-binding residues can result in LKB1 activation.

### Chemical optimization of SMALS

With the identification of the PYP scaffold, we then undertook a modeling-based / structure-based drug design (SBDD) effort to identify a second compound series. Exploring other chemical scaffolds that might occupy the ATP-binding pocket of STRAD and establish similar contacts to NVP-AEW541, we identified the aminopyrazinamide scaffold (APA), with the initial compound SMALS-197 demonstrating activity as the basis of a potential second series (Fig. 2A).

**Figure 2.**
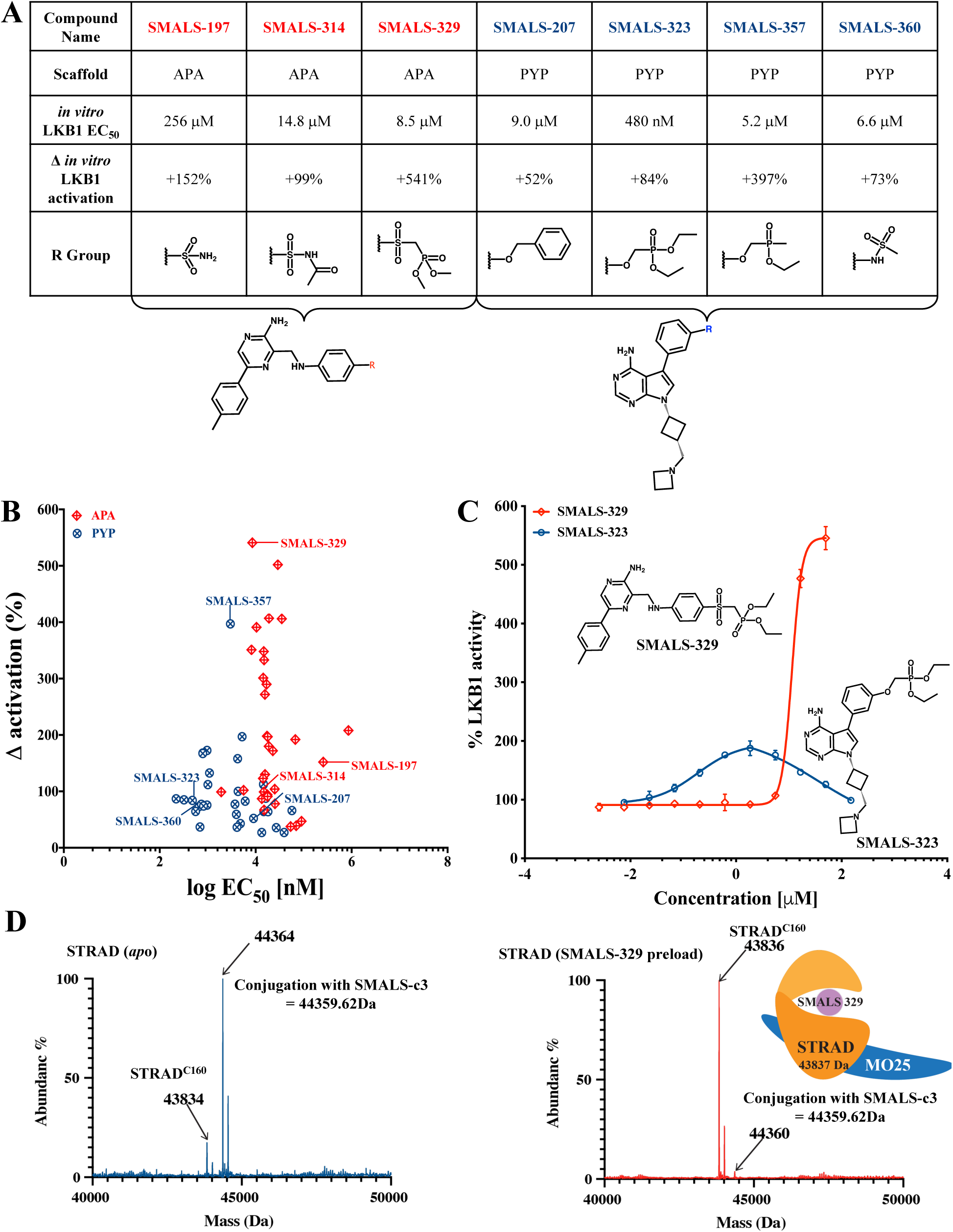
Development of two SMALS scaffolds. A. Selected examples of aminopyrazinamide (APA) and pyrrolopyrimidine (PYP) analogue structure-activity relationship. B. Summary graph of compound potency vs. enhancement of LKB1 activity *in vitro* for APA and PYP scaffolds. PYP compounds had lower EC_50_ values, APA compounds had a greater enhancement of LKB1 kinase activity. C. Dose-response curve for SMAL-323 (PYP) and SMALS-329 (APA), demonstrating differences in ceiling of activation and inhibition of LKB1 at higher concentrations by SMALS-323. D. Competition assay between reversible compound SMALS-329 with irreversible covalent SMALS-c3: Left panel shows SMALS-c3 conjugation to STRAD, which is reduced when STRAD/MO25 is preloaded with 20 µM SMALS-329 for 30 minutes.

Next, we undertook parallel structure activity relationships (SAR) efforts to improve the affinity and activity induction of both scaffolds. We advanced APA and PYP compounds by adding polar functionality to simulate phosphates from ATP, based on the observation from the STRAD crystal structure that 3 polar or positively charged residues directly interact with phosphates in ATP (Lys197, His200, and Arg215; Fig S1A), in place of the magnesium-mediated interactions seen in kinase active sites^35^. These chemical modifications were anticipated to enhance potency while also increasing specificity for STRAD over kinases. Thus, we introduced sulfonamides and ethyl/methyl phosphonates at various positions in this region (Fig 2A; Fig. S1A, B, C).

Our analysis revealed several intriguing differences between these two series: whereas PYP compounds showed higher affinity, they tended to activate LKB1 to a much lesser extent in *in vitro* kinase assays. Conversely our APA series had a relative floor for affinity but was able to increase kinase activity far more substantially than PYP (Fig. 2B). Respective dose-response curves for our APA series lead SMALS-329 (relative EC_50_: 8.5 µM; activation 541%) and our PYP series lead SMALS-323 (relative EC_50_: 480 nM; activation 84%) are shown. A hook effect was also seen at higher doses of SMALS-323 and other PYP compounds (Fig. 2C). In order to evaluate the selectivity of APA and PYP series lead compounds, we also undertook a limited off-target analysis of these compounds and confirmed that they do not significantly impact a set of common off-targets including both LKB1 and IGF1R (Fig. S2).

As our SAR had been driven by an *in vitro* kinase assay, we next directly assessed the binding of each compound to the STRAD/MO25 dimer. As the PYP series was derived from an ATP-displacement assay, it was anticipated that more advanced compounds would still be able to displace ATP from the STRAD/MO25 dimer. Consistent with this, SMALS-323 also displaced TNP-ATP from STRAD with an affinity slightly lower than that of ATP (Fig. S1D).

SMALS-329 and other compounds of the APA series absorb and emit in the same spectra as TNP-ATP. Therefore, we were not able to directly assay ATP displacement by this compound series. Instead, we used SMALS-c3 as a tool compound for the competitive binding with SMALS-329. LC-MS data shows that preloading with SMALS-329 substantially reduces the conjugation with SMALS-c3, suggesting that SMALS-329 binds the same region in STRAD (Fig. 2 D). In addition, preloading with ATP substantially reduces the conjugation with SMALS-c3 supporting our premise that SMALS-c3 binds the ATP binding pocket in STRAD similar to our initial hit NVP-AEW541 (Fig. S1 E). Thus, our data demonstrating that those optimized SMALS compounds bind the ATP pocket. This is consistent with our proposed mechanism of action whereby small molecules binding to the ATP pocket of STRAD activates LKB1.

### SMALS-329 activates LKB1 in tumor cell lines in a target- and dose-dependent manner

We next assessed SMALS effects in cancer cell models. Given that SMALS-329 had the greatest impact on LKB1’s *in vitro* catalytic activity, we subsequently focused on its effect on the LKB1/STRAD/MO25 trimer and the *in vivo* cellular response. We first assessed the direct impact of SMALS-329 on the LKB1 trimer in human tumor cell lines. For this purpose, the A549 *LKB1^Q^*^37^*\**lung cancer cell line was engineered with doxycycline (dox) inducible 3xFLAG-LKB1. Consistent with the known role of LKB1 complex formation in LKB1 activation, we found that SMALS-329 increases the association of LKB1 trimeric complex components measured by immunoprecipitation (Fig. 3A). Similarly, we observed that SMALS-329 not only increases the thermal stability of STRAD by cellular thermal shift assay (CETSA), but also the stability of LKB1 (Fig. 3B).

**Figure 3.**
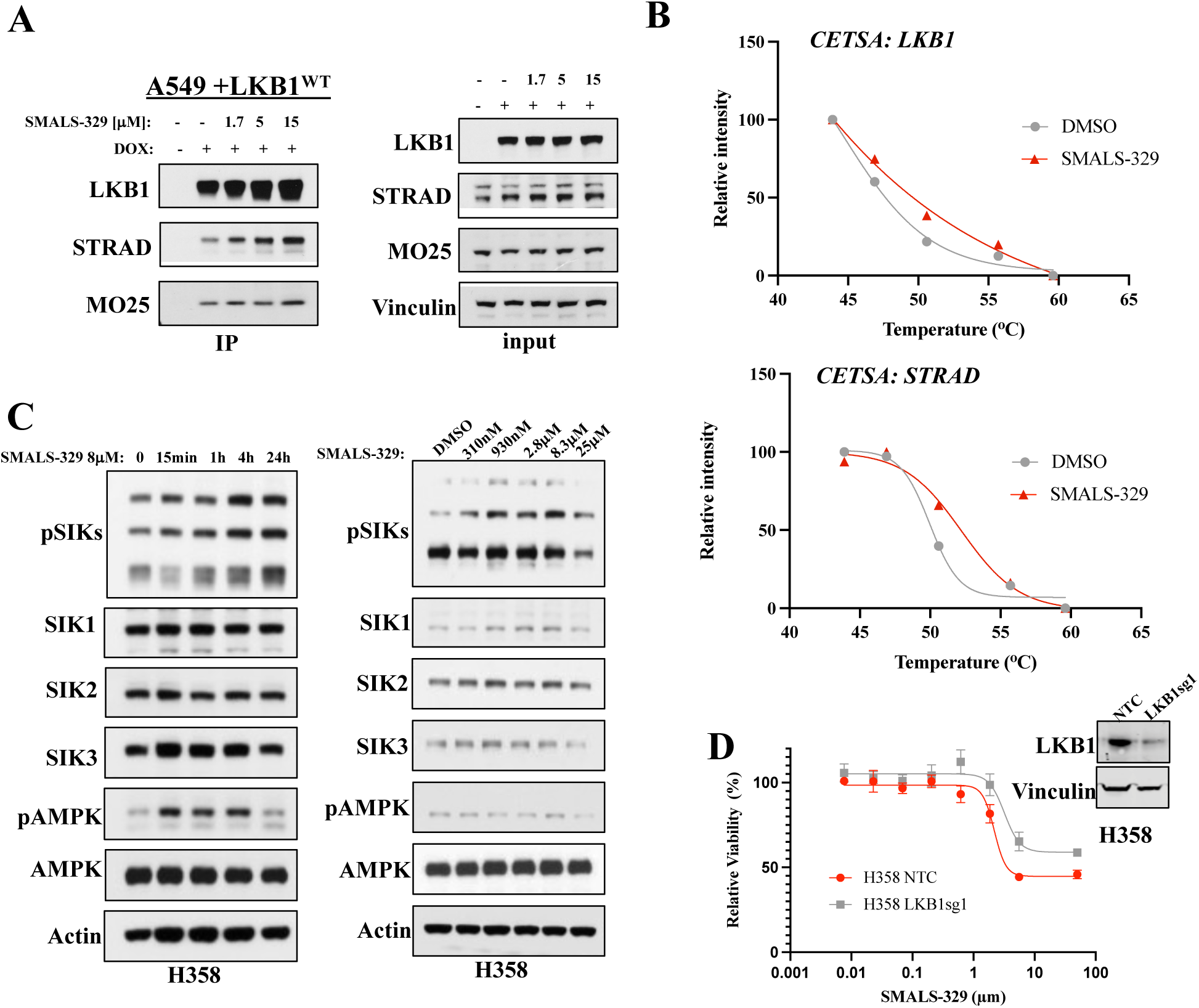
SMALS-329 increase LKB1 trimer association and stability, activating signaling. A. Immunoprecipitation of doxycycline (dox)-inducible 3xFLAG-LKB1^WT^ in A549 cells, a lung cancer line with an early nonsense mutation in *LKB1*. An increase in the association of complex components is correlated with a SMALS-329 dose. B. Quantified results from cellular thermal shift assay in A549 cells. Greater 3xFLAG-LKB1^WT^ and STRAD stabilization is observed following 1 hour treatment with 20 µM SMALS-329 when LKB1^WT^ is stably transduced in A549 cells. C. Left: Western blot analysis of proximal downstream signaling in H358 cells after a time course of 8 µM treatment with SMALS-329. LKB1 activation results in increased SIK phosphorylation. Right: Cells were treated with a range of doses (DMSO, 310 nM, 930 nM, 2.8 µM, 8.33 µM, 25 µM) of SMALS-329 and harvested at 4 hours’ time point. D. Effect of 120 hours of SMALS-329 on cell viability in H358 cells stably transduced with dCas9-KRAB and LKB1 targeting sgRNA or non-targeting control sgRNA, measured by CellTiter-Glo. Inset: western blot confirming knockdown.

To investigate the proximal signaling changes that occur with small molecule stimulation of endogenously expressed LKB1, we used the *LKB1^WT^* H358 lung cancer cell line. We observed that treatment with 8 µM SMALS-329 resulted in increased phosphorylation of the known LKB1 sites on key LKB1 effectors AMPK1/2 and SIK1-3 (Fig. 3C.). Interestingly, SIK phosphorylation peaked at 4 hours and was maintained for 24 hours, whereas AMPK was maximally phosphorylated at 15 minutes and returned to baseline at the 24-hour timepoint. When cells were subjected to doses of SMALS-329 ranging from 0.3 - 25 µM for 4 hours, we observed dose responsive activation up to 8.3 µM, with inhibition at the highest dose. These complex kinetics may be a result of off-target SMALS effects, but also raise the intriguing possibility that distinct cellular feedback processes govern activation of LKB1 effectors.

Next we evaluated the impact of SMALS-329 on lung cancer cell line proliferation. SMALS-329 inhibited the proliferation of H358 cells in a dose-dependent manner, and LKB1 knockdown by CRISPRi reduced this effect by 1.5-fold, suggesting that the inhibitory effect of SMALS-329 depends on expression of its target LKB1 (Fig. 3D). Taken together, we have confirmed that SMALS-329 can engage with the LKB1 complex in cells, activating signal transduction and inhibiting the proliferation of H358 lung cancer cells in a dose- and target-dependent manner.

### SMALS-329 inhibits viability of cell lines derived from multiple tumor types

While the impact of LKB1 inactivation has been deeply studied, the impact of LKB1 activation is much less well understood. Thus, to characterize the broader patterns of cancer cell sensitivity to SMALS, we assembled a panel of 64 cancer cell lines. Given the importance of LKB1 in lung cancer^4^, this panel of tumor cell lines included 32 lung cancer lines. We selected 32 additional cell lines across a broad range of tissue types where LKB1 inactivation has been identified. We used the Cancer Dependency Map ^43^ to guide our selection, focusing on cell lines where LKB1 knockdown or CRISPR deletion resulted in increased cell viability, suggesting that LKB1 activation may in turn reduce cell viability in these cells. We then tested sensitivity to both SMALS-329 and SMALS-357, the PYP compound that induced the highest level of LKB1 activation in an *in vitro* kinase assay (Fig. 2A, B).

12 tumor cell lines showed a GI_50_ ≤ 15 µM for SMALS-329 (Fig.4A, Table. S2). In addition to lung cancer models, endometrial, colorectal, renal and hepatocellular carcinoma models displayed sensitivity to SMALS-329. While SMALS-357 was overall less potent than SMALS-329, we observed a statistically significant positive correlation between PYP and APA compound efficacy, supporting LKB1 activation as the mechanism of growth inhibition. Importantly, none of the cell lines with an *LKB1* inactivating mutation (red dots) showed a GI_50_ <20 µM for either compound (Fig. 4A, Table. S2), corroborating their target dependency. Sensitive and resistant cells represented a wide array of cancers, but neither tissue origin nor tumor types predicted sensitivity to LKB1 activation. We hypothesized that the genomic background of these cells could serve as a better proxy for determining sensitivity. Therefore, we utilized the cancer cell line encyclopedia (CCLE) RNAseq dataset to assess the gene expression-based sensitivity profile^44,45^. We excluded *LKB1* inactivated and genetically *CTNNB1* activated cell lines, given data that LKB1 can directly activate CTNNB1 function^46^. Intriguingly, cancer cell lines resistant to LKB1 stimulation had an enrichment for inflammation relevant hallmark gene sets, including IL-6/JAK/STAT3 signaling and interferon response (Fig. 4B). Upregulation of these gene sets has also been observed in cells with LKB1 loss.

**Figure 4.**
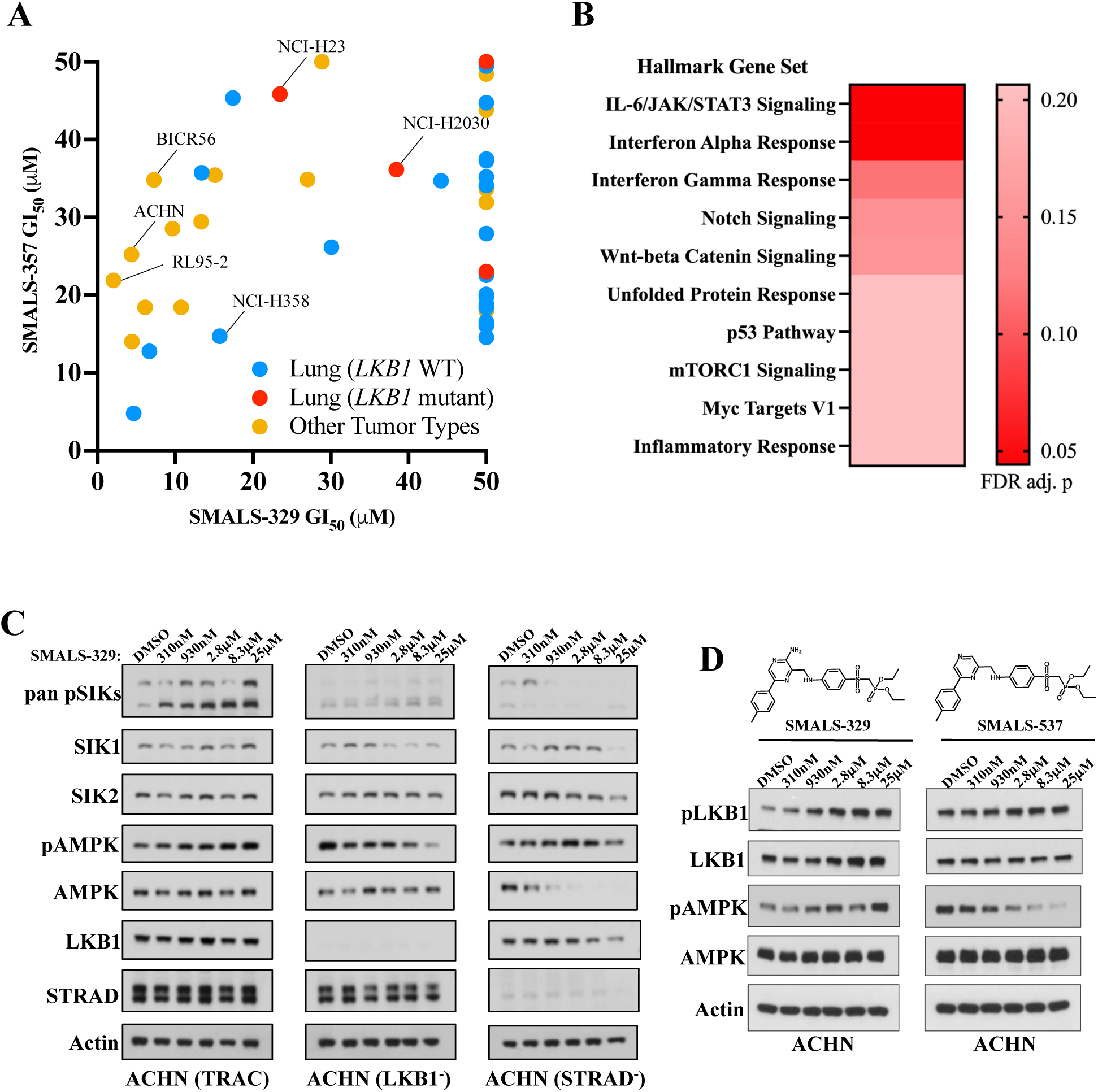
SMAL-329 SMALS compounds effects on growth inhibitor and proximal downstream signal. A. Cell screen to evaluate the growth inhibition of cancer cells by SMALS-329 and SMALS-357 treatment by Cell Titer Glo. Neither tissue of origin nor tumor type predicted sensitivity to LKB1 activation. SMALS-329 GI_50_ vs. SMALS-357 GI_50_: Pearson *r* (62) = .37, *p\*\** = .0026, GI_50_ cut off: 50µM. Four of the most sensitive cells include: ACHN (kidney), BICR56 (tongue), NCI-H358 (lung), and RL95-2 (endometrium). Red dots represent cell lines with LKB1 inactivation. B. Summary gene set enrichment analysis (GSEA) of RNA-seq datasets derived from the Cancer Cell Line Encyclopedia (CCLE) to assess gene expression-based SMALS-329 sensitivity profile. Significant ‘Hallmark’ gene sets are shown. Cell lines sensitive to LKB1 stimulation had downregulation of inflammation-relevant Hallmark Gene Sets. C. Western blot analysis of proximal downstream signaling in ACHN cells with CRISPR deletions of *LKB1*, *STRAD* or T Cell Receptor Alpha Constant (*TRAC*, a CRISPR control). Cells were treated with a range of doses of SMALS-329 (DMSO, 310 nM, 930 nM, 2.8 µM, 8. 3 µM, 25 µM). Treatment with SMALS-329 in control cells resulted in increased phosphorylation of SIKs and AMPK. This effect is lost in cells with CRISPR deletions of *LKB1* and *STRAD*. D. Western blot analysis of proximal downstream signaling in ACHN cells treated with a range of doses of SMALS-329 and a range of doses of a negative control compound lacking the suspected hinge binding motif, SMALS-357. Treatment with SMALS-329 results in increased phosphorylation of LKB1 and increased phosphorylation of AMPK. Treatment with SMALS-537 results in no change in LKB1 phosphorylation and decreased phosphorylation of AMPK.

To further understand LKB1 stimulation in SMALS-329 sensitive cell lines, we treated ACHN and RL95-2 cells with 8 µM SMALS-329 at varying time points (Fig. S3A). We consistently observed the activation of one or several LKB1 effectors with variable signaling kinetics. As showed examples of H358, ACHN, RL95-2 in (Fig.3C, S3A), some tumor cell lines exhibited rapid activation and subsequent downregulation, while others showed prolonged signaling durations.

We used two additional strategies to confirm the specificity of our compounds *in vivo*. While we observed dose-dependent activation of LKB1 signaling by SMALS-329, indicated by the phosphorylation of its substrate sites on AMPK and SIK1-3 in the presence of LKB1 and STRAD, we observed abrogation of the signaling effects driven by SMALS-329 when LKB1 or STRAD were eliminated from ACHN cell via CRISPR knock out (Fig. 4C). This confirmed that the effects of SMALS-329 on downstream signaling are dependent on LKB1 and STRAD. Secondly, we developed a SMALS-329 analogue lacking the exocyclic amine on the pyrazinamide moiety, designated SMALS-537, and found that unlike SMALS-329 this agent did not activate AMPK or LKB1 phosphorylation (Fig. 4D).

Finally, to understand the transcriptional impact of SMAL-329, we performed RNAseq in ACHN cells. Gene set enrichment analysis identified an enrichment of genes involved in cell adhesion/mobility (11 genes) and cell polarity (6 genes) as the key downregulated group of genes. Additionally, we noted significant regulation, both up and down, of several inflammatory mediators and components of the unfolded protein response (UPR (Fig.S3B)). These findings are consistent with the known effects of LKB1, namely the regulation of metabolism, inflammation, cell adhesion/mobility and cell polarity and the UPR.

### SMALS-329 enhances PI3K/AKT inhibitor effects

As LKB1 loss is well known to cause drug resistance, we investigated if augmented LKB1 signaling might potentiate the effects of oncogene inhibition by combining SMALS-329 with a library of kinase inhibitors in ACHN cells. SMALS-329 was tested at 5 µM and kinase inhibitors at 20 nM, 200 nM and 2 µM. Similar results were observed at all 3 concentrations of kinase inhibitors. Notably, the growth inhibition by multiple inhibitors targeting the PI3K/AKT pathway was markedly augmented by SMALS-329. Conversely, a subset of kinase inhibitors was less potent when combined with SMALS-329, most notably the GSK inhibitor CHIR-98014 (Fig. 5A). No significant difference was observed for RAS/MAPK-targeting agents (Fig. 5A). To further evaluate the relationship between SMALS-329 and PI3K/AKT pathway inhibition, we measured dose response curves of the AKT inhibitor GSK2141795, in ACHN, BICR56, and H358 cells. Consistent with our screening data, we observed additive effects with SMALS-329 (Fig. 5B). Cotreatment with SMALS-329 caused a significant decrease in the EC_50_ of GSK2141795 (Fig. 5B, C). As LKB1 is known to impact PI3K/AKT signaling^25^, and both pathways share downstream signaling components, these results support an on-target mechanism of action for SMALS-329. They also demonstrate cooperation between LKB1 activation and oncogenic kinase inhibition in controlling cell proliferation (Fig. 5D).

**Figure 5.**
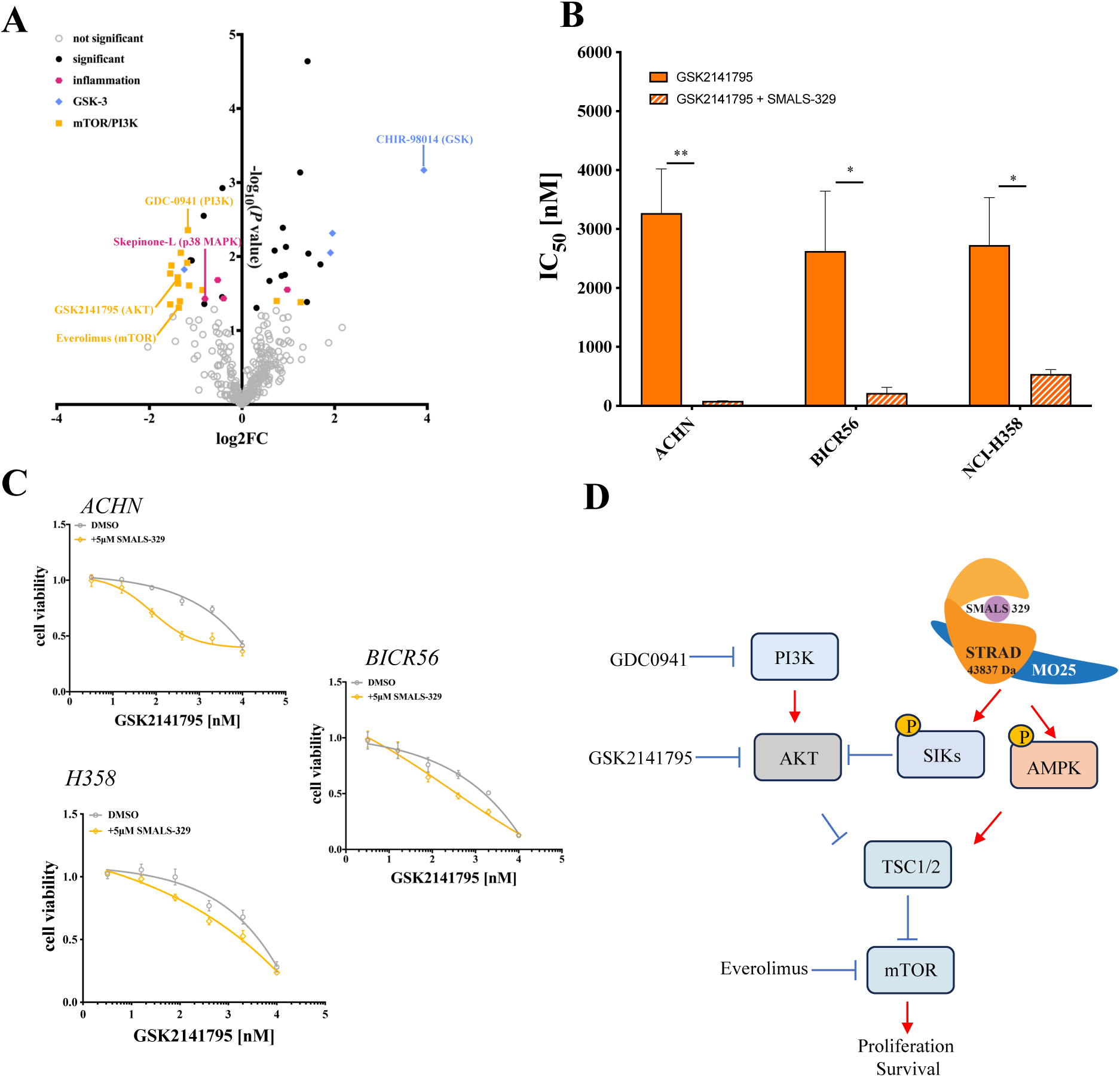
LKB1 activation enhancing the PI3K/AKT inhibitor effects. A. Combination drug screen in ACHN: Cells were treated with 200 nM of kinase inhibitors from the SelleckChem kinase inhibitor library purchased in 2017 and 5 µM SMALS-329. Inhibitors of the PI3K/mTOR pathway components synergized with treatment of SMALS-329. B. Composite Cell Titer-Glo viability assays in ACHN cells. Cells were treated with a dose curve (top dose 10 µM with 1:5 dilutions) of GSK2141795 (AKTi) in combination with DMSO or 5 µM SMALS-329. The addition of SMALS-329 attenuated the effects of these kinase inhibitors, especially with the AKT inhibitor, ** for *p* ≤ 0.01,* for *p* ≤ 0.05. C. Summary graph of the IC_50_ values of Cell Titer Glo experiments in multiple sensitive cell lines treated with GSK2141795 with or without SMALS-329. The addition of SMALS-329 significantly reduced the IC_50_ value of the AKT inhibitor in cells. D. Schematic of proposed mechanism of action.

## Discussion

Here, we report the development of small molecule activators of the tumor suppressor kinase LKB1. These compounds engage the ATP-binding pocket of STRAD and are suspected to act through the allosteric stabilization of LKB1 by STRAD to increase its kinase activity, thereby inhibiting cancer cell lines proliferation and triggering downstream signaling pathways. This report provides proof of concept for augmentation of the function of a non-mutant tumor suppressor kinase as an anti-cancer therapeutic strategy. Tumor suppressor loss has only rarely been a target for precision medicine, with the targeting of synthetic lethal vulnerabilities as a key exception when tumor suppressor function impacts cellular fitness. Several emerging strategies restore the function of a lost tumor suppressor, most notably the development of cysteine-targeted stabilizers of mutant TP53^47^.

Targeting the STRAD pseudokinase with an activator strategy also highlights the opportunity to use allosteric stabilization as a therapeutic modality. Numerous efforts have targeted the ATP binding pocket of oncogenic pseudokinases such as kinase suppressor of Ras (KSR)^48^ and Erb-B2 receptor tyrosine kinase 3 (ERBB3)^49^, and the tyrosine kinase 2 (TYK2) inhibitor deucravacitinib has been approved for psoriasis. Similar to Janus Kinase 2 (JAK2), TYK2 has tandem pseudokinase and kinase domains, with the conformation of the pseudokinase domain influencing kinase activity. Deucravacitinib and the tool compounds above all act on the target protein’s pseudokinase domain by either destabilizing it or favoring an inactive structure^48–50^. Our demonstration that targeting a pseudokinase to generate an active structure suggests a streamlined method of activating a number of other pseudokinase containing proteins and complexes, including the mixed lineage kinase domain-like protein (MLKL), tribbles pseudokinase 3, mixed lineage kinase domain-like (MLKL), Integrin-linked kinase (ILK), PEAK1, kinase suppressor of Ras 1 (KSR1), and general control nonderepressible 2 (EIF2AK4).

Our work reveals challenges unique to development of kinase activators targeting pseudokinase, some of which limit the conclusion that can be drawn from our analysis. Of particular note is the possibility of off target binding of the LKB1 ATP binding pocket. Given this possibility, we have focused our development strategy on engineering specificity in features of STRAD such as direct binding of the phosphates in ATP, resulting in the inclusion of phosphonate and other polar residues. These features did favor specificity, but also limited drug solubility and thus its potential for *in vivo* studies. Future optimization of these compounds is needed to advance pre-clinical development.

Further, while targeting an intact tumor suppressor kinase suggests that these agents could be used in a broad range of cancers, validating therapeutic mechanism of action and identifying predictors of cellular sensitivity will require additional study. Assessment of expression data sets for the cells in our screening panel showed downregulated inflammatory transcription as a marker of SMALS sensitivity, suggesting that these cells might have basal LKB1 activation. Further, while this work focuses on single agent efficacy to inhibit cellular proliferation, drug combination and targeting the TME represent appealing strategies for the use of LKB1 activators. Strong clinical data emerging from non-small cell lung cancer (NSCLC) suggest that LKB1 deficient tumors are resistant to anti-cancer immunotherapy. Thus, better understanding of how SMALS might reverse an immune excluded phenotype in cancer could enable their use to augment immunotherapy. Our work demonstrates the possibility of tackling these exciting questions in cancer therapeutics through the reporting of a tool compound that can directly activate the tumor suppressive function of LKB1.

## Materials and Methods

### Western blot analysis

Cultured cells were washed with PBS and lysed in RIPA lysis buffer (Pierce) with complete protease inhibitor (Sigma Aldrich) and phosphatase inhibitor (Roche). Clarified cell lysates were quantified through BCA assay according to the manufacturer’s instructions (Pierce). Equal amounts of lysate extracts were boiled in 5x loading dye and loaded onto a 4-12% Bis-Tris gel (Invitrogen) and transferred to nitrocellulose membranes (using a standard wet electroblotting system (Bio-Rad)). Membranes were blocked in 5% non-fat dry milk + 0.02% sodium azide in TBS-T, and proteins of interest were analyzed using primary antibodies (1:1000 in 5% BSA in TBS-T) as follows: anti-LKB1 (Cell Signaling Technology; 3050S), anti-phospho-LKB1 (Cell Signaling Technology; 3482S), anti-STRAD (Abcam; ab192879), anti-MO25 (Cell Signaling Technology; 2716S), anti-phospho-AMPK (Cell Signaling Technology; 2535L), anti-AMPK (Cell Signaling Technology; 5832S), anti-pan-phospho-SIKs (Abcam; ab199474), anti-SIK1 (EMD Millipore; ABE799), anti-SIK2 (Cell Signaling Technology; 6919S), anti-SIK3 (Cell Signaling Technology; 39477S), anti-β-actin (Cell Signaling Technology; 3700S), anti-vinculin (Cell Signaling Technology; 13901S). Secondary antibodies used were anti-rabbit IgG, HRP-linked Antibody (Cell Signaling Technology; 7074S) or anti-mouse IgG, HRP-linked Antibody (Cell Signaling Technology; 7076S). Signal intensities were detected using ECL substrate (Pierce) and developed using film. Alternatively, Immunoblots were imaged using the Li-Cor Odyssey CLx Imaging System with Goat Anti-Mouse IgG Antibody, IRDye® 800CW (LI-COR Biosciences; 926-32210), or Goat Anti-Rabbit IgG Antibody, IRDye® 800CW (LI-COR Biosciences; 926-32211) as secondary antibodies (1:20,000 in TBS-T).

### Mammalian cell culture and stable cell line generation

A549 (ATCC) and NCI-H358 (ATCC) cells were cultured in RPMI-1640 (Corning) supplemented with 10% FBS (Life Technologies) and 1% penicillin/streptomycin (Gibco) at 37°C and 5% CO_2_. ACHN (ATCC) cells were cultured in MEM (Corning) supplemented with 10% FBS (Life Technologies) and 1% penicillin/streptomycin (Gibco) at 37°C and 5% CO_2_. RL95-2 (ATCC) cells were cultured in DMEM: F12 (Gibco) supplemented with 10% FBS (Life Technologies), 1% penicillin/streptomycin (Gibco), and 0.005mg/mL insulin (Gibco) at 37°C and 5% CO_2_. All cells were occasionally tested for mycoplasma contamination and confirmed by STR if not purchased. For lentivirus production, plasmids were packaged in HEK 293T cells for 72 hours using lipid-based transfection. The N’-terminal 3x FLAG A549 LKB1^WT^, NCI-358 LKB1 knock down, NCI-H358 NTC cell lines were generated through lentiviral transduction followed by selection with puromycin (Gibco).

### Protein expression and purification

His-STRAD^C160^ and MO25 or His-STRAD^S160^ and MO25 were co-expressed in E. coli BL21 using the pOPCH vector. Wild-type STRAD^C160^ and MO25: His-STRAD (residues 59-431) and MO25 (residues 1-341) sequences were obtained from Arvin Dar’s lab and codon optimized for bacterial expression. Site-directed mutagenesis was employed to mutate the STRAD residue C160 to S160. The PCR reaction was set up with a final concentration of 1X Q5 Hot Start High-Fidelity Master Mix (New England Biolabs). The forward and reverse primers were used at a final concentration of 0.5 μM each. The sequences of the primers were as follows: Forward Primer: ACA GCT GAT TAG CAC TCA CTT CAT G, Reverse Primer: TTC GCG GAG CCG TAC. KLD For the KLD reaction, 1 μl of the PCR product was mixed with 1X KLD Reaction Buffer and 1X KLD Enzyme Mix (New England Biolabs). The reaction mix was then used to transform BL21 chemically competent cells. The sequence of mutated STRAD^S160^and MO25: His-STRAD (residues 59-431 with C160S mutation) and MO25 (residues 1-341) was confirmed by whole plasmid sequencing performed by Primordium.

The purification procedure was similar to the protocol outlined in^35,36^, but modifications were made to accommodate the available lab equipment. Bacteria were induced for 18 h with 250 μM IPTG, pelleted and resuspended in the lysis Buffer (50 mM Tris pH 7.8, 50 mM NaCl, 10% glycerol, 20 mM imidazole, 0.2 mM EGTA, 0.2 mM EDTA, 0.075% β-mercaptoethanol,0.5 mg/mL lysozyme (ThermoFisher; 89833), and 0.3 mg/mL Benzonase (Millipore Sigma; E1014)). Cells were lysed using a microfluidizer for 2 cycles. Lysate was clarified with ultracentrifugation at 30,000 x g for 30 min and loaded onto Ni-NTA resin (Qiagen; 30210). After two washes in the Lysis Buffer and one in the Lysis Buffer with 40 mM imidazole, protein was then eluted in Lysis Buffer with 200 mM imidazole. The supernatant was dialyzed overnight at 4°C using Slide-A-Lyzer™ Dialysis Cassettes (Thermo Fisher Scientific). The sample was then filtered through a 0.22-µm filter and further purified by size-exclusion chromatography (SEC) using a Superdex 75 Increase 10/300 GL column (Cytiva). Fractions were collected on an FPLC, with the STRAD-MO25 dimer eluting early while STRAD monomer eluted later. Purity was then confirmed by SDS-PAGE and Coomassie staining to assess contaminants and separate STRAD monomer from STRAD-MO25 dimer-containing fractions.

### LC-MS analysis of covalent conjugation and competing binding

Wild-type STRAD^C160^–MO25 or mutation STRAD^S160^–MO25 at 2 μM was reacted with SMALS-c3 at 20 μM (2% (v/v) dimethylsulphoxide (DMSO) final) in 25mM Tris-Hcl, 150 mM NaCl on rocker. The conjugation was assessed by electrospray mass spectrometry using a Waters Acquity UPLC/ESI-TQD with a 2.1 × 50 mm Acquity UPLC BEH300 C4 column at time point 6 hours. Spectra were deconvoluted and analyzed using Masslynx V4.2 software. For competing binding, Wild-type STRAD^C160^–MO25 was pre-incubated with reversible compound SMALS-329 or ATP at 20 μM for 30mins on a rocker. The sample was then reacted with SMALS-c3 at 20μM. The competitive binding was analyzed by LC-MS as described above.

### Cell viability assays

Cells were seeded into 96-well tissue culture treated plates at 500 cells per well and incubated at 37°C and 5% CO_2_ overnight. Cells were treated with selected drugs at different final concentrations and incubated for another 120 hours. After incubation, plates and CellTiter-Glo® (Promega) reagent were allowed to equilibrate at room temperature on the bench for 30 min. Luminescence was measured on a SpectraMax M3 plate reader or BioTek Synergy 2 plate reader to quantify the cell viability.

Similarly, for the cancer cell line screen assay. A panel of 64 cancer cell lines were treated with a range of doses of SMALS-329 or SMALS-357 and luminescence measured using the CellTiter-Glo assay. ACHN cell line combination drug screen was measured with CellTiter-Glo. Cells were treated with 200nM of kinase inhibitors from the SelleckChem kinase inhibitor library and 5µM SMALS-329. Combination treatment assay in ACHN, BI3R56, and H358was measured by Cell Titer Glo. Cells were treated with a range of doses (top dose 10µM with 1:5 dilutions) of GSK2141795 (AKTi) in combination with DMSO or 5µM SMALS-329. Data were analyzed with GraphPad Prism software.

### Immunoprecipitation

A549 LKB1^WT^ cells were incubated with doxycycline for 24h at 37°C and incubated with a dose curve of SMALS-329 or DMSO for 1h at 37°C. Cells were washed with PBS and lysed in low salt buffer with complete protease inhibitor (Sigma-Aldrich) and phosphatase inhibitor (Roche). To immunoprecipitate the LKB1-MO25-STRAD complex, magnetic anti-FLAG beads (Millipore Signma) were used according to manufacturer’s instructions. Cleared lysates were incubated with rotation for 2.5h at 4°C with washed anti-FLAG beads. Following immunoprecipitation, the anti-FLAG beads were washed three times using low salt buffer (50mM HEPES pH 7.5, 150mM NaCl, 0.5% Triton X-100, 1mM EDTA). Samples were eluted with 5 μ L of 5x loading dye and analyzed by western blotting. anti-LKB1 (1:1000 in 5% BSA in TBS-T; Cell Signaling Technology; 3050S), anti-STRAD (1:1000 in 5% BSA in TBS-T; Abcam; ab192879), anti-MO25 (1:1000 in 5% BSA in TBS-T; Cell Signaling Technology; 2716S)

### Cellular thermal shift assay (CETSA)

A549 LKB^WT^ cells were incubated with doxycycline for 48h at 37°C and then incubated with 20μM SMALS-329, or DMSO for 1h at 37°C. Scraped cells were resuspended in PBS containing protease and phosphatase inhibitors before being heated for 3min at various temperatures. Cells were snap frozen in a dry ice and ethanol bath and lysed by repeated freeze thaw cycle. Cell lysates were cleared by centrifugation to remove cell debris as well as precipitated and aggregated proteins. Protein expression levels were determined by western blotting.

### TNP-ATP assay

A buffer solution was prepared consisting of 50 mM Tris (pH 7.5), 50 mM NaCl, and 1 mM DTT. STRAD-MO25 was prepared at a concentration of 5 μM. TNP-ATP was also prepared at a concentration of 5 μM. STRAD-MO25 and TNP-ATP were pre-incubated in the prepared buffer solution for 10 minutes at room temperature. Following pre-incubation, compounds were added to the mixture at a concentration of 50 μM. After an additional 10 minutes incubation at room temperature, fluorescence was measured using a spectrofluorometer with an excitation wavelength of 410 nm and an emission wavelength of 538 nm.

### *In vitro* LKB1 activation assay

The *in vitro* LKB1 activation assay was conducted in 384-well plates with a Caliper device. The detailed protocol is available as reported earlier^42^. The 5x kinase assay buffer comprised 200 mM MOPS-NaOH (pH 7.0), 5 mM EDTA, 10 mM DTT, 50 mM MgCl2, and 0.5% Triton X-100. For the assay solutions, the compound solution per well included 1x kinase assay buffer (final), DMSO or compound, ddH_2_O was added to make a final volume of 5 μL. The enzyme solution per well consisted of 1x kinase assay buffer (final), 5 nM LKB1-STRA-/MO25 trimer (Sigma-Aldrich,14-596), ddH_2_O was added to make a final volume of 10 μL.The substrate solution per well contained 1x kinase assay buffer (final), 2.5 μM Fam-labeled LKBtide (Anaspec, AS-64871), 50 μM ATP, ddH_2_O was added to a final volume of 10 μL. 5 μLof the compound assay solution was added to each well, followed by 10 μL of the enzyme solution and incubating for 15 minutes at room temperature. Subsequently, 10 μL of the substrate/ATP solution was added, and the mixture was incubated for 2 hours at room temperature. The reaction was terminated by adding stop buffer (150 mM EDTA). Samples were analyzed using a Caliper device with settings for sample buffer (100 mM HEPES, pH 7.5, 1 mM EDTA, 0.015% Brij-35, 0.1% CR-3, 5% DMSO), upstream voltage of -270 V, downstream voltage of -1850 V, pressure of -1.2 psi, sample sip time of 0.3 s, and buffer sip time of 90 s. The retention times were 225–230 s for the product and 240 s for the substrate. Percent conversion (% conversion) was calculated using the equation: % conversion = Prod / (Prod + Subst). Data were plotted using GraphPad Prism software, and EC50 values were determined using a four parameters nonlinear fit. Fold activation was calculated by comparing the % conversion with drug to the baseline.

### Eurofins selectivity profiling

Kinase selectivity profiling was performed using the KINOMEscan™ platform (Eurofins Discovery) with 97 targets tested. The assay uses a competition-based binding approach, where the test compound (10μM) competes with an immobilized ATP-site-directed ligand. Bound kinases were detected using a streptavidin-tagged detection system and measured by ultra-sensitive qPCR, and the results were expressed as percentage of control binding (%Ctrl). Processors were described previously in^51,52^.

### RNAseq

Following established protocols in the Ramani lab^53^. Total RNA was extracted from ACHN cells using the Total RNA Purification Kit (Norgen Biotek), following the manufacturer’s instructions. Reverse transcription was performed using SuperScript II Reverse Transcriptase (Thermo Fisher Scientific). Library preparation was conducted by the Cornell Transcriptional Regulation and Expression (TREx) Facility using the NEBNext Ultra II Directional Library Prep Kit (New England Biolabs). Sequencing was performed on the Illumina NextSeq500 platform at the Genomics Facility of the Cornell University Biotechnology Resource Center, generating paired-end reads with a minimum of 80 M reads per sample. Reads were aligned to the human genome, and normalization and differential gene expression analyses were performed using DESeq2. Each sample achieved at least 25 million uniquely mapped reads, with a unique mapping rate exceeding 90%.

### Small Molecule Synthesis

All compound synthesis was performed at Chempartner (Shanghai). Detailed synthetic methods are available in US patent WO2021155004^54^.

## Author contributions

Lin Song Kretschmer: Conceptualization, Data curation, Formal analysis, Investigation, Visualization, Methodology, Writing – original draft, Writing – review and editing.

Mitchell Dominique: Conceptualization, Data curation, Formal analysis, Investigation, Visualization, Methodology, Writing, – original draft.

John Gordan: Supervision, Conceptualization, Data curation, Formal analysis, Investigation, Visualization, Methodology, Resources, Funding acquisition, Writing – original draft, Writing – review and editing.

LeeAnn L. Wang: Data curation, Formal analysis, Investigation

Jonathan M. L. Ostrem: Conceptualization, Methodology, Covalent compound design, Writing – review and editing.

Rigney E. Turnham: Investigation, Writing – review and editing.

Yeonjoo C. Hwang, Weicheng Li, Aidan Keith, Vijay Ramani, Timothy Kellett, Gary K. L.Chan: Investigation

Roopa Ramamoorthi: Alliance Management and project discussions, Writing – review and editing. Kliment A. Verba: Conceptualization, Methodology

Jin Liu, Zheng Wang, Changliang He, Luo Ding, Tingting Qing, Marc Adler, Sarah Lively, Richard Beresis: Conceptualization, Investigation, Methodology

Robert Drakas: Conceptualization, Resources, Funding acquisition, Methodology

## Acknowledgments

We are grateful to Kevan Shokat, Natalia Jura and Arvin Dar for their insight and advising of this project and for the strong support of Michael Xin Hui throughout this effort.

We appreciate the contributions of Shanghai Yingli Pharmaceutical for their synthesis of several covalent compounds (including SMALS-c3).

We thank Fujimori lab and Shokat lab for providing access to LC-MS, we thank Qinheng Zheng, Darius Mcardle, and Patrick Pfaff for their assistance with LC-MS. We thank Ostrem lab, Kortemme lab and Verb lab for providing access to microfluidizer, ultracentrifuge, FPLC, protein purification relevant chemical and consumables. We thank Giuseppe Cimicata, Lieza Chan, Yilun Qi for their assistance with the protein purification process. We thank Rohit Bhadoria for providing chemical support. We thank Rony Francois for discussions. We thank Simon Kretschmer for manuscript editing and scientific advising.

We thank lab members of Gordan, Ostrem and Korgan labs for helpful discussions, feedback and advice throughout this project.

## Data and materials availability

All data are available in the main text or the supplementary materials. Correspondence and requests for materials should be addressed to John Gordan

## Supplementary Figures and Tables

**Supplementary Figure 1.**
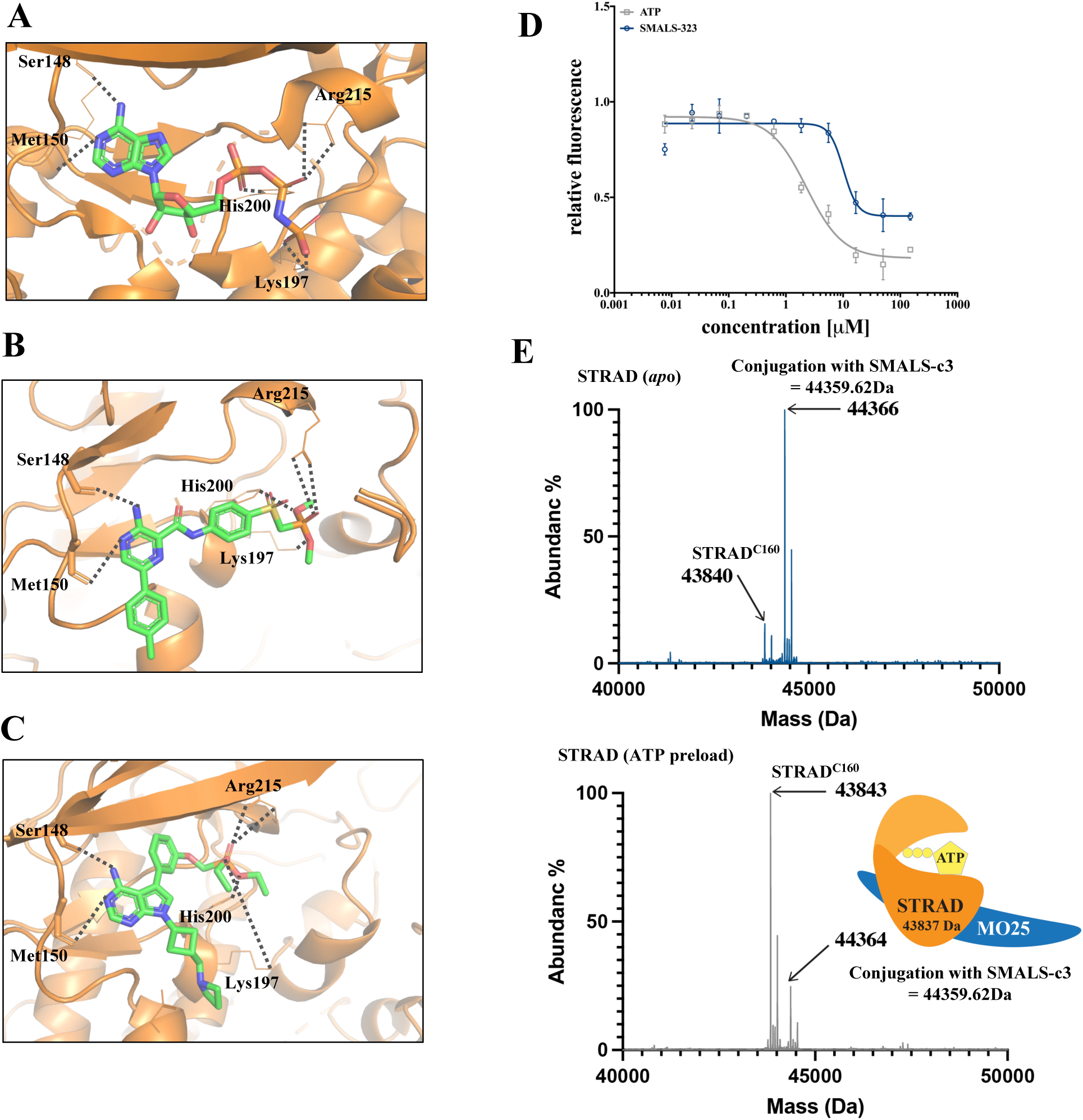
Development of a second class of SMALS. A. Crystal structure of ADP-PNP in STRAD’s ATP-binding pocket from 2WTK. Ser148 and Met150 coordinate with the adenosine aromatic ring for hinge binding. Lys 197, His200, and Arg215 coordinate with the phosphate groups. B. PYMOL computational modeling of aminopyrazinamide (APA) scaffold in STRAD’s ATP-binding pocket, proposed contacts shared with ATP are shown. C. PYMOL computational modeling of PYP scaffold in STRAD’s ATP-binding pocket with proposed contacts. D. TNP-ATP displacement by ATP and SMALS-323 in recombinant dimeric STRAD/MO25. E. Competition assay between ATP with irreversible covalent SMALS-c3: Upper panel shows SMALS-c3 conjugation to STRAD, which is reduced when STRAD/MO25 is preloaded with 20 µM ATP for 30 minutes.

**Supplementary Figure 2.**
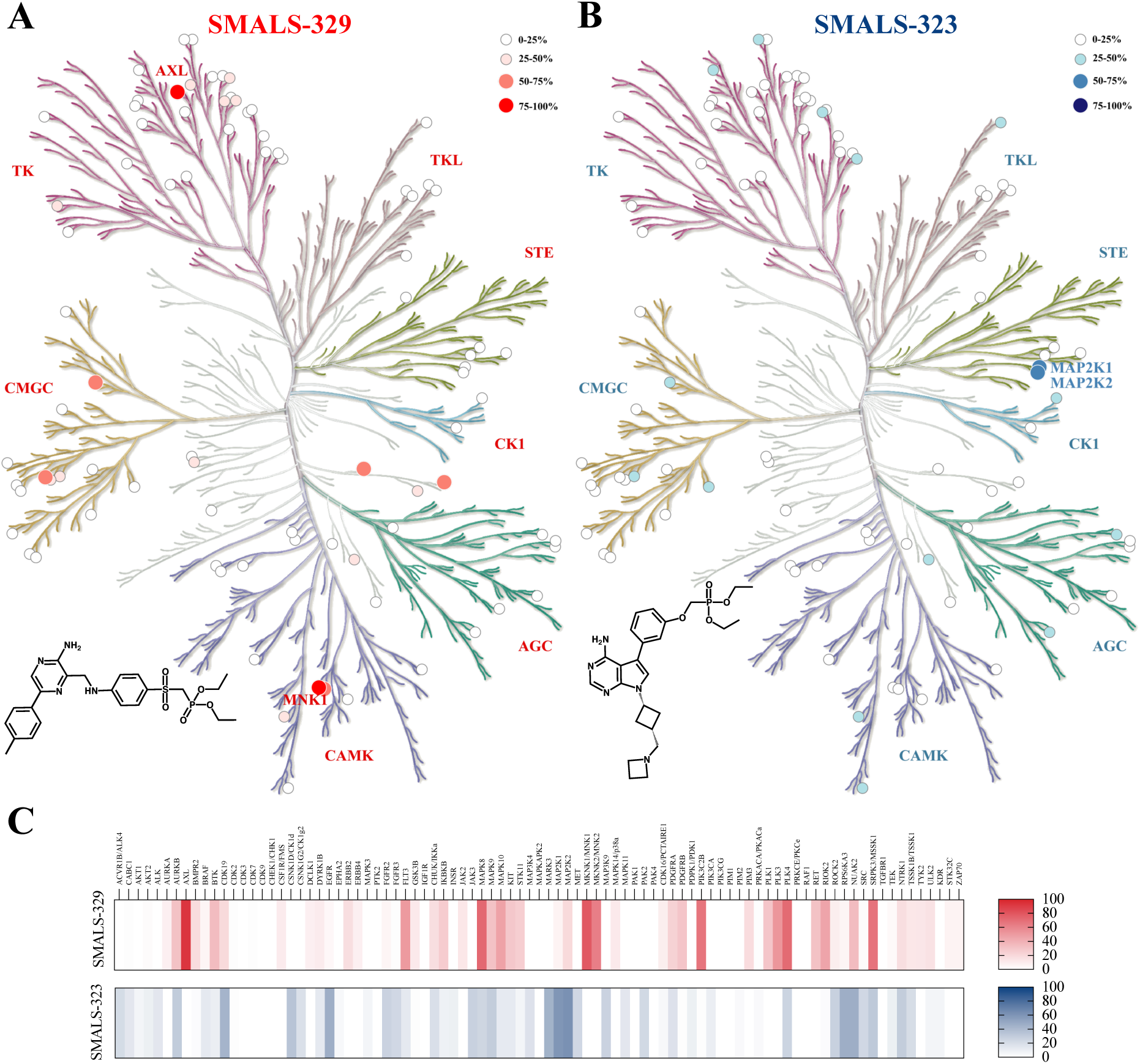
Development and comparison of a second class of compound able to bind STRAD to activate LKB1. A. Eurofins off-target kinome profiling of SMALS-329. SMALS-329 had 6 moderate off-target hits of 50-75% inhibition: FLT3, PLK3, MNK2, MSSK1, PLK4, and JNK1. SMALS-329 had 2 strong off-target hits of 75-100% inhibition: MNK1 and AXL. B. Eurofins off-target kinome profiling of SMALS-323. SMALS-323 had 2 moderate off-target hits of 50-75% inhibition: MAP2K1 and MAP2K2. SMALS-323 had no strong off-target hits of 75-100% inhibition. C. Heatmap of off-target kinome profiles of hit compounds SMALS-329 and SMALS-323, showing very little overlap or similarities between off-targets of each compound.

**Supplementary Figure 3.**
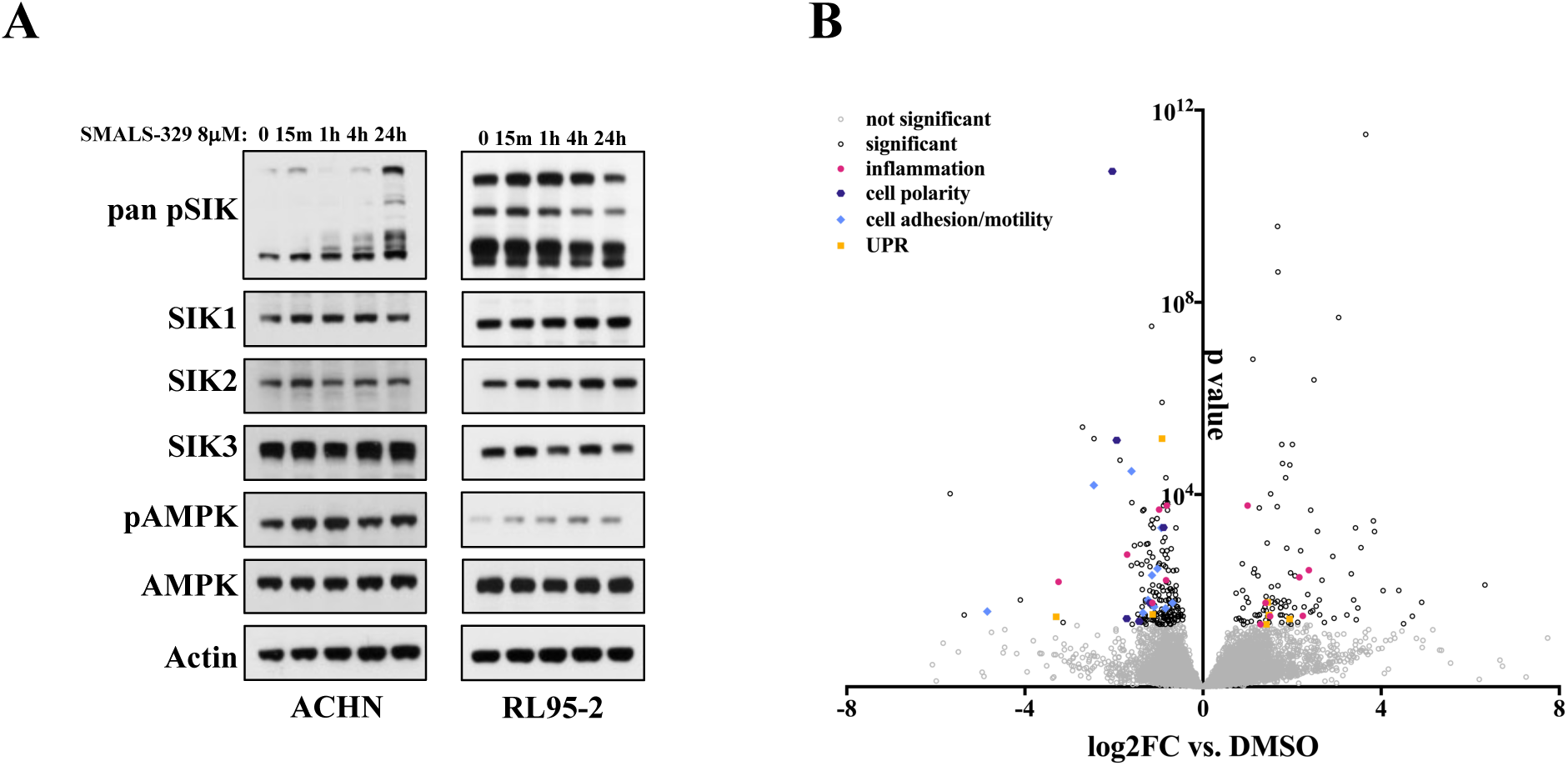
Signaling and transcriptional effects of SMALS-329. A. Western blot analysis of proximal downstream signaling in ACHN and RL95-2 after a time course of 8 µM treatment with SMALS-329. LKB1 activation results in increased SIKs and AMPK phosphorylation, albeit with distinct kinetics. B. RNAseq analysis in ACHN after 24h treatment with 8 µM SMALS-329. Hits regulating cell polarity (all on downregulated side), cell adhesion/motility (all on the downregulated side), inflammation and unfolded protein response (UPR) are highlighted.

**Supplementary Table 1.**
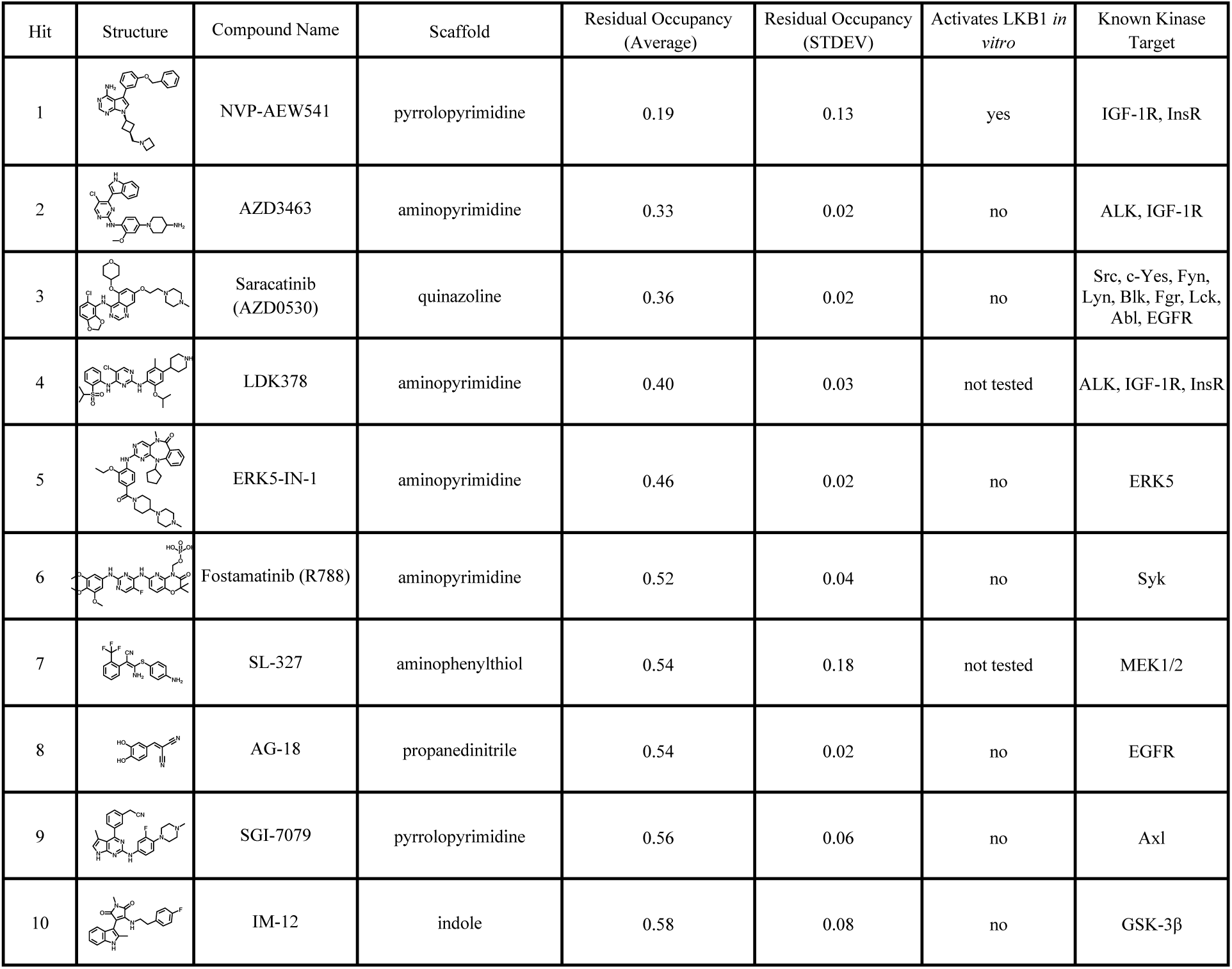
Table of top 10 hits of TNP-ATP screen showing compound structure, compound name, average residual occupancy ±std dev, ability to activate LKB1 *in vitro*, and known kinase target. The compound with the lowest residual occupancy and greatest ability to bind STRAD’s ATP-binding pocket was NVP-AEW541, an IGF-IR inhibitor with a PYP scaffold.

| Hit | Structure | Compound Name | Scaffold | Residual Occupancy (Average) | Residual Occupancy (STDEV) | Activates LKB1 *in vitro* | Known Kinase Target |
| --- | --- | --- | --- | --- | --- | --- | --- |
| 1 |  | NVP-AEW541 | pyrrolopyrimidine | 0.19 | 0.13 | yes | IGF-1R, InsR |
| 2 |  | AZD3463 | aminopyrimidine | 0.33 | 0.02 | no | ALK, IGF-1R |
| 3 |  | Saracatinib (AZD0530) | quinazoline | 0.36 | 0.02 | no | Src, c-Yes, Fyn, Lyn, Blk, Fgr, Lck, Abl, EGFR |
| 4 |  | LDK378 | aminopyrimidine | 0.40 | 0.03 | not tested | ALK, IGF-1R, InsR |
| 5 |  | ERK5-IN-1 | aminopyrimidine | 0.46 | 0.02 | no | ERK5 |
| 6 |  | Fostamatinib (R788) | aminopyrimidine | 0.52 | 0.04 | no | Syk |
| 7 |  | SL-327 | aminophenylthiol | 0.54 | 0.18 | not tested | MEK1/2 |
| 8 |  | AG-18 | propanedinitrile | 0.54 | 0.02 | no | EGFR |
| 9 |  | SGI-7079 | pyrrolopyrimidine | 0.56 | 0.06 | no | Axl |
| 10 | | IM-12 | indole | 0.58 | 0.08 | no | GSK-3β |

**Supplementary Table 2.** Cell screen cell lines. Table of the 64 cell lines (including 32 lung cancer lines) used in our cell screen. Listed are the cell line, tissue of origin, type of cancer, and the relative GI_50_ of compounds SMALS-329 and SMALS-357 respectively. Neither the tissue of origin nor tumor type correlates with the response to LKB1 activation. All LKB1 inactivated cell lines are resistant lines. Red: STK11 frameshift, nonsense, or mutations result in a truncated and inactive protein product and occur most commonly located within the kinase domain and classified as likely oncogenic based on OncoKB™^55^. Cell line signaling mutation information was retrieved from CCLE 2019 report when available^56^; otherwise, from the CCLE 2012 report^45^ using cBioPortal.

| Cell Line | Tissue | Tumor Type | SU-329 GI50 (μM) | SU-357 GI50 (μM) | Oncogenetic signaling Mutations |
| --- | --- | --- | --- | --- | --- |
| NCI-H209 | lung | small cell | 4.64 | 4.79 | TP53 X225_splice; RB1 C706F |
| Saos-2 | bone | osteosarcoma | 4.42 | 14.02 | RB1 X605_splice |
| NCI-H1299 | lung | non-small cell | 6.67 | 12.77 | NRAS Q61K |
| LS513 | colon | adenocarcinoma | 6.11 | 18.44 | KRAS G12D; SMARCA4 R973W;RAF1 E478K |
| RL95-2 | endometrium | carcinoma | 2.05 | 21.9 | PTEN R173H; HRAS Q61H; PTEN M134I; PIK3R1 R638*; SOS1N233I;TP53 V218del |
| NCI-H358 | lung | non-small cell, bronchioalveolar carcinoma | 15.73 | 14.72 | KRAS G12C |
| HT-1080 | connective tissue | fibrosarcoma | 10.76 | 18.42 | NRAS Q61K |
| ACHN | kidney | adenocarcinoma | 4.38 | 25.21 | MET AMP; EGFR AMP; KRAS AMP |
| OV7 | ovary | carcinoma | 9.65 | 28.55 | KRAS G12D; SMARCA4 Q1185*; TP53 I254S |
| EBC-1 | lung | squamous carcinoma | 50 | 14.57 | TP53 E171* |
| ChaGo-K-1 | lung | bronchogenic carcinoma | 50 | 16.03 | TP53 C275F; TP53 X187_splice |
| SNU398 | liver | carcinoma | 13.33 | 29.43 | CTNNB1 S37C; TP53 X224_splice; SMARCA2 R1246* |
| A427 | lung | carcinoma | 50 | 20.09 | KRAS G12D; CTNNB1 T41A |
| BICR 56 | tongue | squamous cell carcinoma | 7.26 | 34.8035 | NOTCH1 E216*;TP53 X126_splice |
| NCI-H441 | lung | papillary adenocarcinoma | 30.1 | 26.17 | KRAS G12V;TP53 R158L |
| MCF-7 | breast | adenocarcinoma (pleural effusion) | 50 | 18.06 | PIK3CA E545K |
| OVTOKO | ovary | clear cell adenocarcinoma | 15.11 | 35.42 | PIK3R1 N453dup |
| Calu-6 | lung | anaplastic carcinoma | 13.38 | 35.72 | KRAS Q61K;TP53 R196* |
| BICR 22 | tongue | squamous carcinoma | 27 | 34.885 | PIK3CB A1048V; PTPN11 G503V; TP53 X307_Splice; tp53 F109Gfs*37 |
| NCI-H526 | lung | variant small cell | 50 | 7.04 | TP53 X33_splice |
| Calu-1 | lung | epidermoid carcinoma | 50 | 27.9 | KRAS G12C |
| NCI-H1975 | lung | non-small cell | 44.2 | 34.7 | EGFR T790M; EGFR L858R; PIK3CA G118D |
| HCC44 | lung | non-small cell | 50 | 23.02 | KRASG12C; **STK11 M511fs\*112 FSins** |
| NCI-H2052 | lung | mesothelioma (pleural effusion) | 50 | 18.4 | MYC AMP |
| NCI-H1581 | lung | non-small cell | 50 | 18.83 | TP53 I144* |
| NCI-H2030 | lung | non-small cell | 38.435 | 36.14 | KRASG12C; **STK11 E317\* nonsense mutation** |
| NCI-H226 | lung | squamous cell carcinoma | 17.4 | 45.36 | MYC AMP |
| NCI-H23 | lung | non-small cell | 23.45 | 45.85 | KRASG12C; **STK11 W332\* nonsense mutation** |
| NCI-H661 | lung | large cell | 50 | 16.57 | SMARCA4 L1161Sfs*3; TP53 R158L; TP53 S215I |
| MSTO-211H | lung | biphasic mesothelioma | 50 | 22.85 | N/A |
| NCI-H2228 | lung | non-small cell | 50 | 19.72 | TP53 Q331* ; RB1 X204_splice |
| OE33 | esophagus | carcinoma | 50 | 17.99 | TP53 C135Y |
| HPAF-II | pancreas | adenocarcinoma | 50 | 33.63 | KRAS G12D; TP53 P151S |
| YAPC | pancreas | carcinoma | 28.87 | 50 | KRAS G12V; TP53 H179R |
| NCI-H292 | lung | carcinoma | 50 | 22.57 | N/A |
| 786-O | kidney | carcinoma | 50 | 50 | BRAF V600E; NOTCH2 M1? |
| COLO 829 | skin | melanoma | 50 | 43.82 | BRAF V600E |
| ASPC1 | pancreas | adenocarcinoma | 50 | 50 | KRAS G12D; TP53 C135Afs*35 |
| OVSAHO | ovary | adenocarcinoma | 50 | 22.73 | TP53 R342* |
| HCC15 | lung | non-small cell | 50 | 44.75 | KRAS AMP; MYC AMP |
| BT-483 | breast | ductal carcinoma | 50 | 50 | PIK3CA E542K; NF1 R1204W; TP53 M246I |
| CAL33 | tongue | squamous cell carcinoma | 50 | 50 | PIK3CA H1047R; TP53 R175H |
| NCI-H322 | lung | bronchioalveolar carcinoma | 50 | 34.1 | KRAS AMP |
| NCI-H1155 | lung | non-small cell | 50 | 37.52 | MYCN AMP |
| CAPAN1 | pancreas | adenocarcinoma | 50 | 50 | KRAS G12V; TP53 A159V |
| NCI-H1650 | lung | bronchoalveolar carcinoma | 50 | 37.28 | EGFR E746_A750del; TP53 X225_splice |
| MKN7 | stomach | adenocarcinoma | 50 | 48.44 | NF1 X547_splice; TP53 I251L; RB1 X905_splice |
| PC-9 | lung | adenocarcinoma | 50 | 31.92 | EGFR E746_A750del; TP53 R248Q; NOTCH3 R1893* |
| NCI-H650 | lung | non-small cell | 50 | 35.2 | KRAS Q61L; TP53 K164N |
| Hs 294T | skin | melanoma | 50 | 50 | BRAF V600E; TP53 R280S |
| Hs 766T | pancreas | carcinoma | 50 | 50 | KRAS Q61H; |
| NCI-H508 | colon | adenocarcinoma | 50 | 49.46 | PIK3CA E545K; BRAF G596R |
| NCI-H1648 | lung | adenocarcinoma | 50 | 50 | TP53 L35Ffs*8 |
| Panc 10.05 | pancreas | adenocarcinoma | 50 | 50 | KRASG12D; TP53 I255N |
| RMUG-S | ovary | carcinoma | 50 | 50 | TP53 A347V |
| SCC-4 | tongue | squamous cell carcinoma | 50 | 50 | TP53 P151S |
| SK-MEL-2 | skin | melanoma | 50 | 50 | NRASQ61R; TP53 G145S |
| SNU423 | liver | carcinoma | 50 | 50 | TP53 X126_splice; KEAP1 X511_splice |
| YD-8 | tongue | squamous cell carcinoma | 50 | 50 | N/A |
| SW1573 | lung | alveolar cell carcinoma | 50 | 50 | KRAS G12C; PIK3CA K11E; SMARCB1 X121_splice |
| A549 | lung | carcinoma | 50 | 50 | KRAS G12S; KEAP1 G333C; **STK11 Q37\*nonsense mutation** |
| NCI-H2122 | lung | adenocarcinoma | 50 | 50 | KRAS G12C; TP53 C176F; **STK11 P281Rfs\*6 (FS del)** |
| PLC/PRF/5 | liver | hepatoma | 50 | 50 | SMARCA4 K1332Sfs*26; TP53 R249S; TP53 G199*; **STK11 D194N** ( on kinase domain) |
| H727 | lung | Carcinoid | 50 | 50 | KRAS G12V; TP53 Y163_Q165DUP; STK11 Q302P |

